# Unraveling Molecular Mechanisms of Low-Salinity Adaptation in *Fenneropenaeus chinensis*: Insights from Gill Tissue Transcriptomics

**DOI:** 10.1101/2020.11.12.379511

**Authors:** Jun Liu, Rongchen Liu, Hui Jiang, Hai Zhang, Dianyuan Zheng, Daizhen Zhang

## Abstract

This study investigates the molecular responses of Chinese shrimp (*Fenneropenaeus chinensis*) to low salinity stress, employing RNA-seq and functional enrichment analyses. A total of 12,332 expressed genes were identified, with 227 differentially expressed genes (DEGs) detected, revealing intricate adaptive responses in gill tissues. Functional enrichment analysis highlighted DEGs associated with ion-transport regulation, bio-metabolism and detoxification (based on GO terms). The upregulation of *ATP1A1*, *SLC26A2*, and *STK39* suggested a pivotal role of ion-transport regulation, emphasizing the importance of maintaining osmotic balance. In terms of bio-metabolism, *ENPP6*, *GLS*, and *DMGDH* exhibited significant upregulation, indicating adjustments in energy metabolism for low-salinity adaptation. Notably, the downregulation of *GSTO1* and *CTH* raised questions about detoxification pathways, potentially reflecting adaptations or compromises in response to low salinity. Further correlation network analysis highlighted key genes like *GLS*, *STK39*, *SESN1*, and *SARDH*, revealing intricate relationships among DEGs. RT-qPCR validation confirmed the consistency of gene expression patterns with RNA-seq results. In conclusion, this study provides a comprehensive understanding of the molecular mechanisms underlying Chinese shrimp responses to low salinity stress. The identified genes lay the groundwork for targeted interventions and strategies to enhance shrimp aquaculture system resilience amid changing salinity levels.

## 1. Introduction

The adaptation of commercial aquaculture crustaceans to low salinity has garnered significant attention [1–4]. It is necessary to understand their adaptation ability to low salinity and mechanisms used by these organisms to tolerate low-salinity environments on purpose of improving the low-salinity aquaculture of marine crustaceans. As osmoregulators, some euryhaline marin and brackish crustaceans have a strong ability to adapt to environments with varying salinities (from almost 0 ppt up to 40 ppt) [5–8]. Therefore, this ability to tolerate low salinity is a key factor affecting the distribution of such crustaceans in low-salinity environments [9–10].

Previous studies have explored the physiological mechanisms underlying low-salinity tolerance in mariculture crustaceans at the organismal, cellular, and molecular levels [8, 11–14]. In general, the most common adaptive strategies for hyperosmoregulation aim to maintain hemolymph osmolarity above that of the ambient medium, both via salt absorption and via permeability reduction (i.e., reducing or limiting water inflow) [6]. However, most euryhaline crustaceans produce isosmotic urine, and thus considerable salt is lost in low-salinity environments [15–16]. Therefore, osmoregulation should be the focal point of our considerations. Studies of low-salinity tolerance in crustaceans have shown that the gills participate in osmoregulation. In detailed reviews, Péqueux (1995) and Henry (2012) assessed the specialized functions of gills and gill parts in various crustaceans and showed that both cuticle permeability and the membrane characteristics of the salt-transporting gill epithelial cells were critical to osmoregulation [6, 8]. Some bio-molecules in the gill epithelial cells facilitate salt absorption, and these biomolecules include Na^+^/K^+^-ATPase, K^+^ channels, Cl^-^ channels, carbonic anhydrase, aquaporins (AQPs) and various exchangers (Na^+^/NH_4_^+^, Na^+^/H^+^, and Cl^-^/HCO_3_^-^) [6, 9, 17–20]. With development of RNA-seq technology, high-throughput screening of genes in response to low salinity stress in crustaceans is currently feasible.

The euryhaline Chinese shrimp (*Fenneropenaeus chinensis*), which has an isosmotic point of 25 ppt, is naturally distributed primarily in the Chinese Yellow Sea, the Bohai Sea, and along the western coast of the Korean Peninsula [21–22]. It is an important commercial shrimp along the coasts of China and Korea [21]. As these shrimp are cultured in much lower salinity (under the isosmotic point), they must manage or tolerate substantial changes in water osmolality. The poorer ability limits the farming development of *F. chinensis* compared to *Penaeus vannamei* [22]. In our prior investigation, an experimental design was implemented to assess the low-salinity tolerance of the target species. However, the experimental protocol involved a group-wise allocation of three prawns each, designated as the low-salinity group (LS), which were subjected to a salinity level of 5 ppt for a duration of 24 hours. To further investigate the molecular mechanisms underlying the low-salinity adaptation of the euryhaline Chinese shrimp, this study aims to fill a necessary gap in our understanding of how these key physiological processes are orchestrated at the molecular level, particularly within the gill tissue with a more meticulous design.

## 2. Materials and Methods

### 2.1 Sample collection and treatment

From July to August 2020, live shrimp were obtained from a farming pool in Lianyungang (N 34^◦^48′ 52.47″, E 119^◦^12′19.08″) and transported to our laboratory at Lianyungang Normal College. All shrimp were acclimated to a salinity of 25 ppt at the beginning. Shrimp acclimation and water-salinity control were reported in our latest research [23]. In short, gill tissues from a random subset of surviving individuals in each group (six from each) were used for transcriptome analysis and real-time quantitative PCR (RT-qPCR).

### 2.2 RNA isolation, library construction, and sequencing

The method here was consistent with that of our previous work[23]. Total RNA was extracted using TRIzol reagent (Invitrogen Corp., USA). RNA concentration was measured using a NanoDrop 2000 spectrophotometer (Thermo Scientific, USA), and RNA integrity was assessed using 1.5% agarose gel electrophoresis. Magnetic oligo (dT) beads were used to isolate mRNA from total RNA. The mRNA was then fragmented into fragments approximately 200 bp long using fragmentation buffer (Tris-acetate, KOAc, and MgOAc) at 94°C for 35 min. The fragmented mRNA was used to construct the cDNA libraries. At least 5 μl of mRNA solution (≥ 200 ng/μl) was used to construct each library. Sequencing libraries for each sample were generated using the TruSeq RNA Sample Prep Kit (Illumina, USA). The read length was 300bp.

### 2.3 Transcriptome Assembly, Gene Functional Annotation, and Analysis of Differentially Expressed Genes

Prior to assembly, the raw reads were subjected to filtering using fastp (v0.23.4) [24]. Reads containing adapters, exceeding 10% of unknown nucleotides, exhibiting low quality, or originating from rRNA were excluded from further analysis. This filtering step aimed to obtain a set of high-quality clean reads. Subsequently, the clean reads were mapped to the Penaeus chinensis reference genome (ASM1920278v2) using HISAT2.2.1 [25]. The mapped reads were then assembled using StringTie (v1.3.1) [26–27] in a reference-based approach. This process ultimately yielded the reads and TPM (transcripts per million) expression matrix.The identified unigenes were annotated against five databases: NCBI nonredundant protein sequences (NR), Protein Families (PFAM), KEGG (Kyoto Encyclopedia of Genes and Genomes), Gene Ontology (GO), and eggNOG [28–29].

Differential expression analysis of RNAs between the FCN10 and FCN25 groups was conducted using the DESeq2 (v1.6.3) software. Genes exhibiting the false discovery rate (FDR) parameter < 0.05 and |Log2 FC| ≥ 1 were considered differentially expressed genes (DEGs). To evaluate functional annotation and pathway enrichment, Gene Ontology (GO) and Kyoto Encyclopedia of Genes and Genomes (KEGG) analyses were performed on DEGs. Statistically significant GO terms and KEGG pathways with padj < 0.05 were determined via statistical enrichment analysis[23]. Z-score was also used to investigate the significance of GO terms[30]. Correlation analysis entails utilizing the stat package based on R (v4.2.3) and employing the Pearson correlation coefficient to assess the correlation between genes[31].

### 2.4 Verification of DEGs using RT-qPCR

We selected 8 RNA-Seq DEGs (7 upregulated and 1 downregulated) for RT-qPCR validation. We used the *TUBA* (Tubulin) as the internal reference gene, against which to normalize the expression levels of the target genes. It was stably expressed in this study. Gene-specific primers were designed based on sequences derived from the transcriptome assembly and annotation using Primer Premier 5.0 (Table 1). Each RT-qPCR (25 μl) contained 12.5 μl of 2 × SYBR qPCR Mix, 1 μl each of forward and reverse primers, 1 μl of cDNA, and 10.5 μl of RNase-free H2O. RT-qPCRs were performed on an Applied Biosystems 7500 Real-time PCR system (Applied Biosystems, Thermo Fisher Scientific, Waltham, MA, USA), with the following cycling conditions: an initial denaturation step of 3 min at 95°C; 40 cycles of 15 s at 94°C, 15 s at 55°C, and 25 s at 72°C; and a standard dissociation cycle. Three technical replicates were performed per gene, and the 2^−△△CT^ method was used to calculate relative expression levels[32].

**Table 1.**
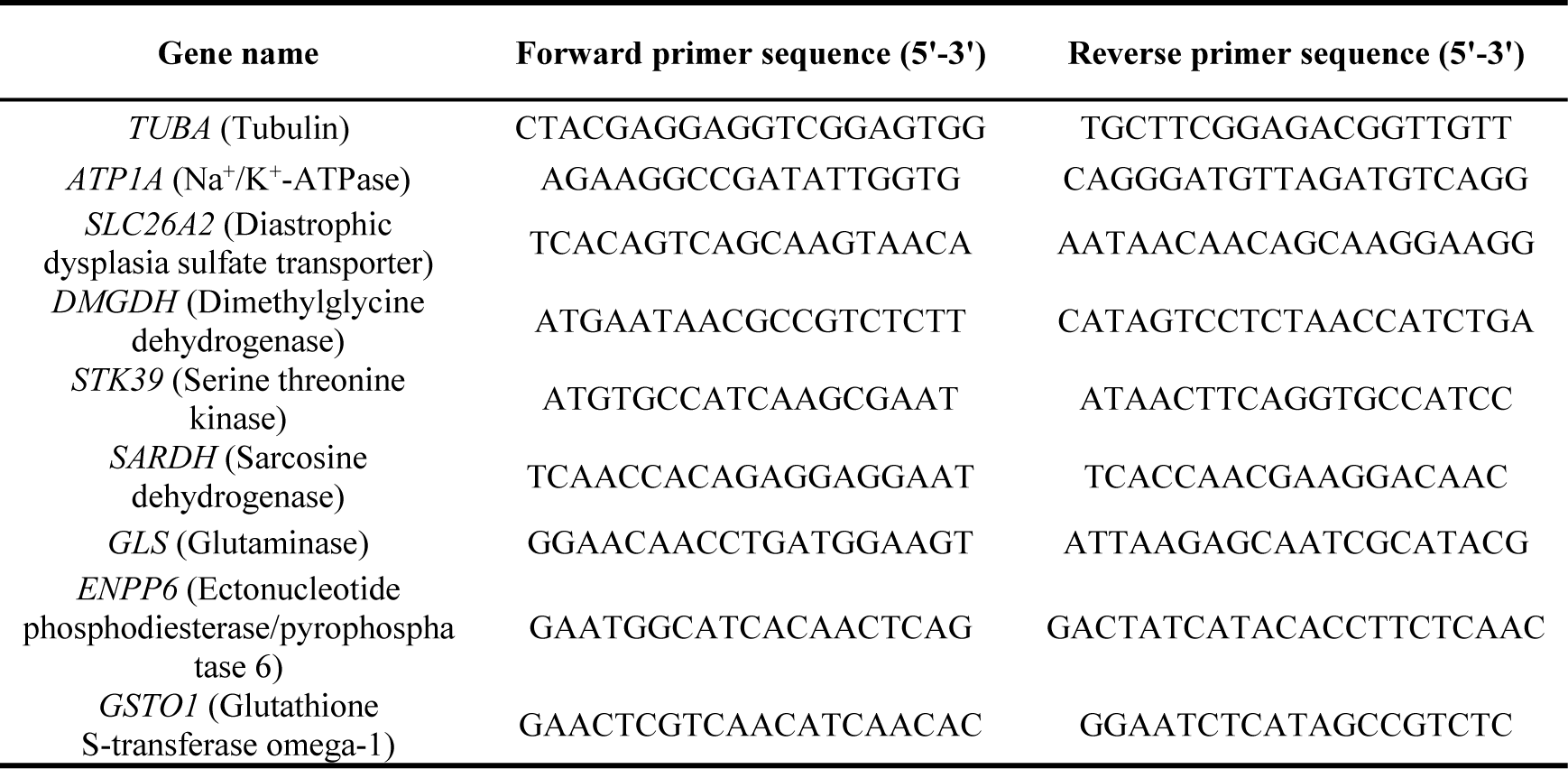
Primers used for real-time quantitative PCR.

## 3. Results

### 3.1 Sequencing and Annotation

After sequencing and raw data cleaning, this research yielded 65.05 Gb data with more than 38 million clean reads for per library. The value of Q30 (bases with quality scores > 30) was more than 90% in average, indicating a high accuracy in base recognition (Table 2). Totally 20, 076 unigenes were annotated against databases of NR, GO, KEGG, eggNOG and PFAM, accounting for 98.9% of unigenes (Table 3).

**Table 2.** Profile of sequencing data.

| Sample | Clean reads | Q30 bases | Alignment rate | Group ID |
| --- | --- | --- | --- | --- |
| SRR14718710 | 38.32M (98.62%) | 5.18G (91.68%) | 87.81% | FC25_1 |
| SRR14718711 | 40.45M (98.57%) | 5.57G (93.07%) | 87.82% | FC25_2 |
| SRR14718712 | 38.32M (98.42%) | 5.24G (92.65%) | 87.24% | FC25_3 |
| SRR17931570 | 56.67M (97.51%) | 7.74G (91.66%) | 88.88% | FC25_4 |
| SRR17931571 | 46.43M (97.38%) | 6.28G (90.89%) | 89.64% | FC25_5 |
| SRR17931572 | 49.36M (97.43%) | 6.75G (91.91%) | 88.40% | FC25_6 |
| SRR14718716 | 40.27M (98.47%) | 5.52G (93.07%) | 85.79% | FC10_1 |
| SRR14718719 | 43.52M (98.74%) | 5.98G (93.00%) | 87.85% | FC10_2 |
| SRR14718720 | 40.17M (99.09%) | 5.55G (93.15%) | 88.56% | FC10_3 |
| SRR17931576 | 52.22M (97.40%) | 7.12G (91.77%) | 86.52% | FC10_4 |
| SRR17931579 | 45.64M (97.50%) | 6.27G (92.14%) | 87.90% | FC10_5 |
| SRR17931580 | 53.78M (97.54%) | 7.35G (91.70%) | 82.92% | FC10_6 |

**Table 3.** Summary of unigenes’ annotation.

| Database | Number of Unigenes | Rate(%) |
| --- | --- | --- |
| NR | 14, 810 | 72.9% |
| PFAM | 7, 302 | 36.0% |
| eggNOG | 7, 398 | 36.5% |
| GO | 6, 279 | 30.9% |
| KEGG | 5, 611 | 27.8% |
| Annotated in at least one database | 2, 0076 | 98.9% |

### 3.2 DEGs in gill of F. chinensis and their functional enrichment

According to the screening results by DESeq2, we detected 12, 242 expressed genes (Supplement 1) and identified 227 DEGs, including 109 up-regulated genes and 118 down-regulated genes (Figure 1, Supplement 2).

**Figure 1.**
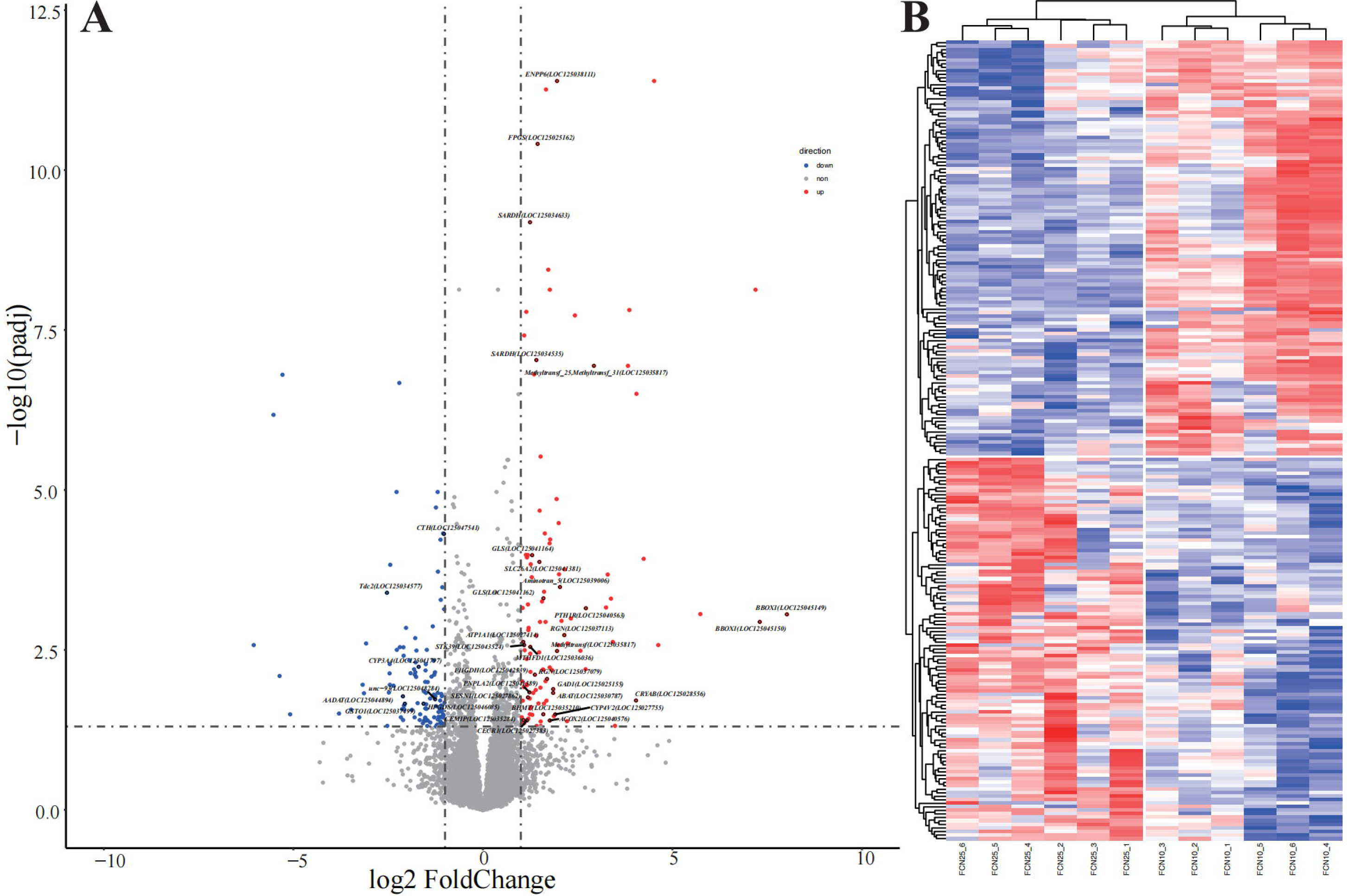
Profiles of gene expression. (**a**) Volcano plot depicting all genes under low salt stress, with up-regulated differently expressed genes represented in red (log2 Fold Change value ≥1) and down-regulated differently expressed genes represented in blue (log2 Fold Change value ≤ −1); (**b**) Selected 227 differently expressed genes in response to low salt stress to construct the heat map.

Further enrichment analysis based on GO showed 120 GO terms were significantly enriched by DEGs (p adjust ≤ 0.05). These GO terms are mainly classified into three categories (Figure 2): the first category is related to ion transport, including “negative regulation of ion transmembrane transport” (GO: 0034766), “negative regulation of ion transport” (GO: 0043271), and “positive regulation of transmembrane transport” (GO: 0034764) etc.; The second category is associated with bio-metabolism, including “serine family amino acid biosynthetic process” (GO: 0009070), “cellular amino acid biosynthetic process” (GO: 0008652) and “vitamin biosynthetic process” (GO: 0009110) etc.; The third category involves processes related to detoxification, including “oxidoreductase activity, acting on the CH-NH group of donors(GO: 0016491); cellular oxidant detoxification (GO: 0098869); cellular detoxification (GO: 1990748) etc. (Description of all GO terms were listed in Supplement 3).

**Figure 2.**
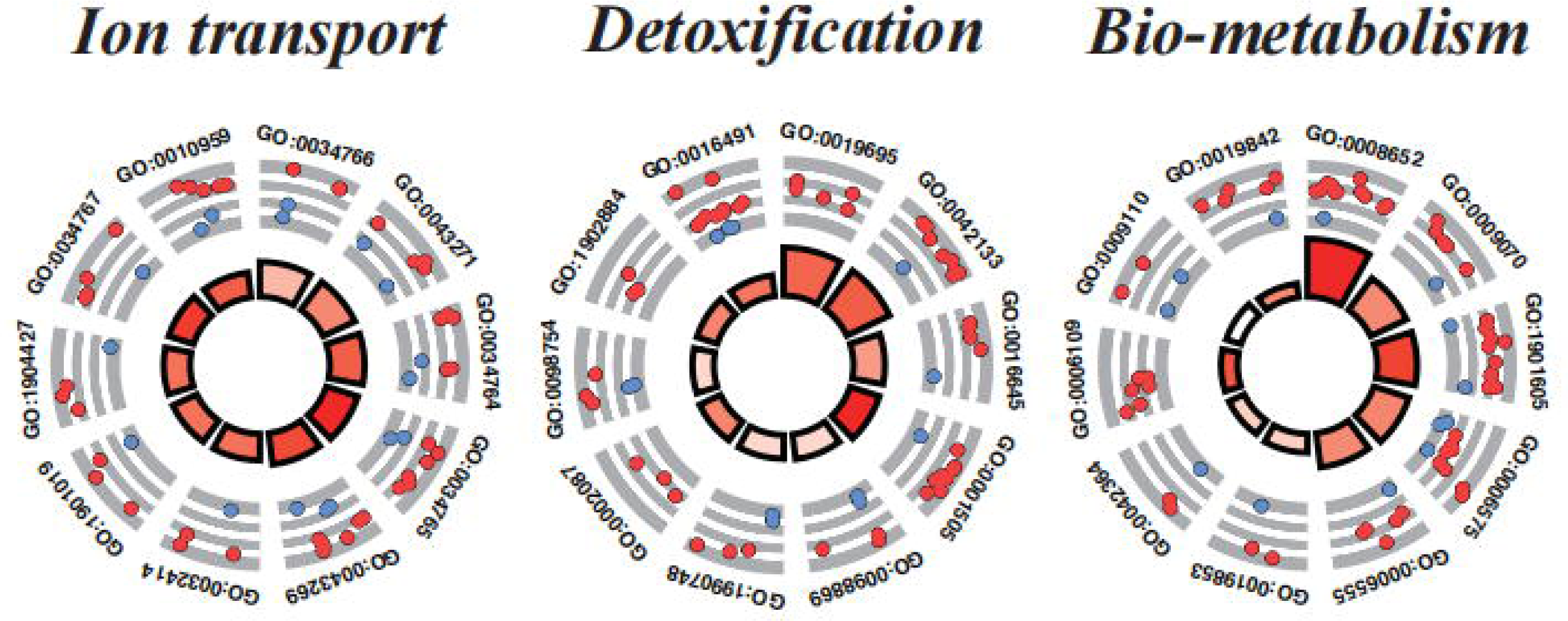
Profiles of gene expression. GO enrichment analysis on DEGs. Three concentric circles represent ion transport, detoxification and bio-metabolism. Each square in the outer circle represents specific GO terms associated with each functional category, with red dots indicating up-regulated genes and blue dots indicating down-regulated genes. The size and color intensity of each red square in the inner circle represent the magnitude of the Z-score.

Results of KEGG enrichment analysis showed DEGs enriched in pathways which are also associated with ion transport, bio-metabolism and detoxification (Figure 3), including “Proximal tubule bicarbonate reclamation”, “Ascorbate and aldarate metabolism”, “GABAergic synapse”, “Alanine, aspartate and glutamate metabolism”, “Protein digestion and absorption” and so on.

**Figure 3.**
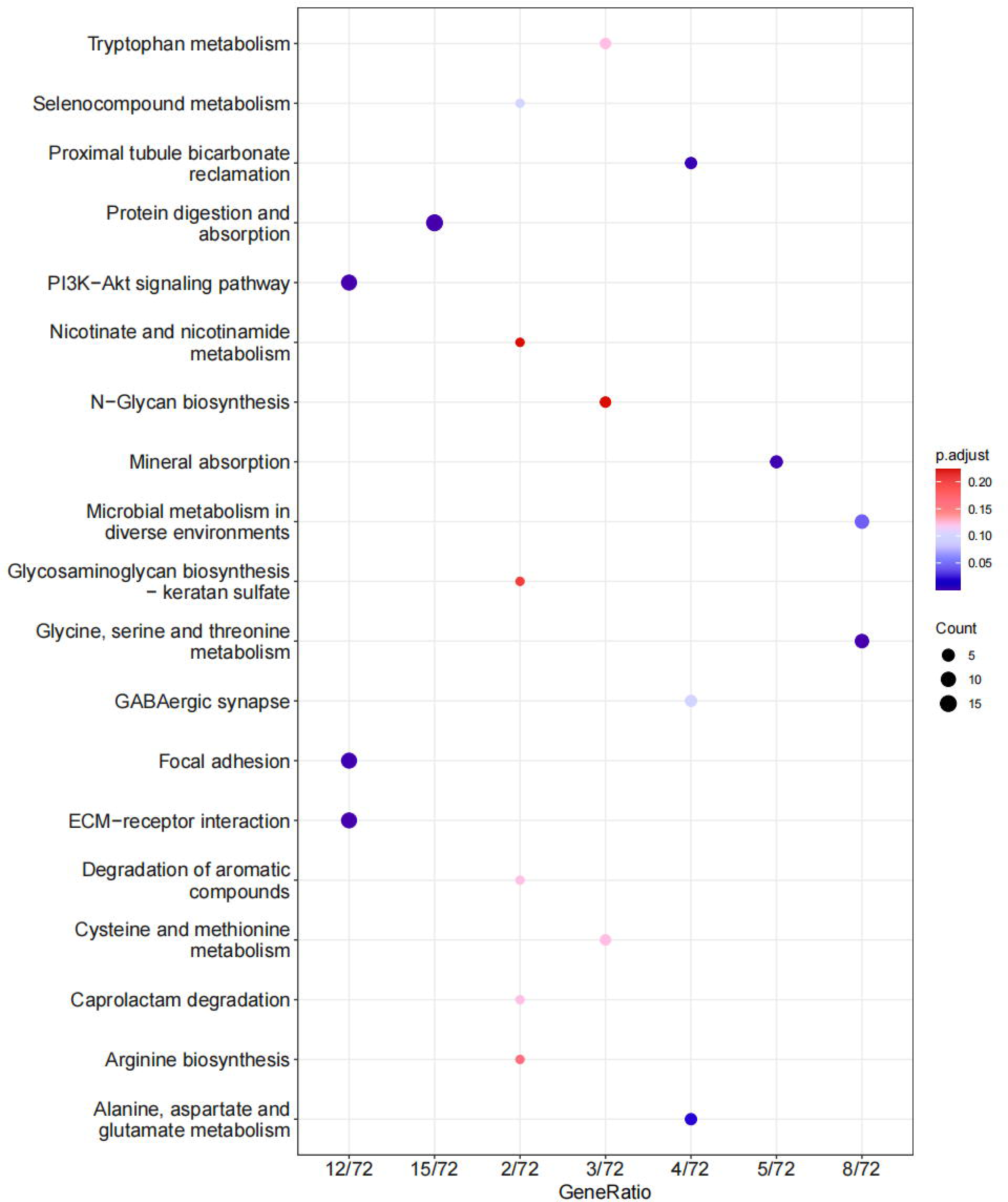
KEGG enrichment analysis on DEGs. This chart illustrates the gene count and the proportion it represents in relation to the total annotated genes for each KEGG metabolic pathway. The y-axis denotes the names of the KEGG metabolic pathways, while the x-axis represents the number of genes.

### 3.3 Correlationship among DEGs

Based on the results of DEGs functional enrichment, a total of 32 DEGs were selected to construct a correlation network, focusing on processes related to ion transport, bio-metabolism and detoxification (Figure 4). The findings revealed a notable correlation pattern among these genes. Specifically, genes such as *GLS* (Glutaminase), *STK39* (Serine threonine kinase), *SESN1* (PA26 p53-induced protein), and *SARDH* (Sarcosine dehydrogenase) exhibited a significantly positive correlation with most of the 32 DEGs, while *CTH* (Cystathionine gamma-lyase) displayed a marked negative correlation with the majority of these DEGs.

**Figure 4.**
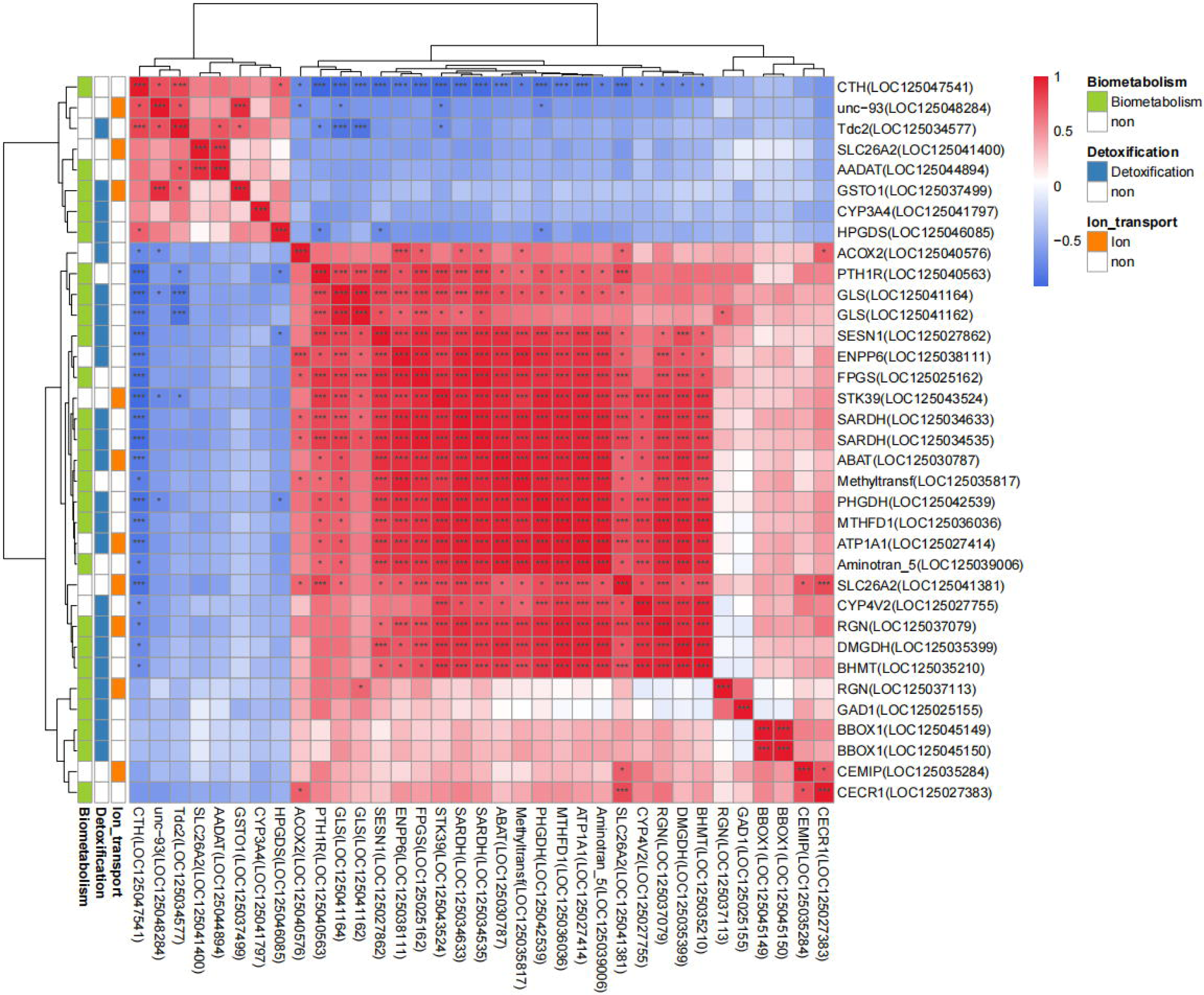
Illustration of pairwise correlation among key DEGs. Heatmap illustrating the pairwise correlation among 32 genes, represented by colors indicating the strength of correlation. Red color means positive relation, Blue color means negatively relation. “*” denotes genes with p-value less than 0.05, “**” represents p-value less than 0.01, and “***” indicates p-value less than 0.001.

### 3.4 RT-qPCR verification

To verify the reliability of the RNA-seq results, we selected 8 representative genes related to low-salinity stress response in aspects of ion-transport regulation, bio-metabolism and detoxification using RT-qPCR method. The results showed that the expression patterns of DEGs were consistent with the RNA-seq results. Specifically, in the low-salinity treatment group, the expression of *ENPP6*, *SARDH*, *SLC26A2*, *GLS*, *ATP1A1*, *STK39* and *DMGDH* genes were up-regulated, while the expression of *GSTO1* gene was down-regulated (Figure 5).

**Figure 5.**
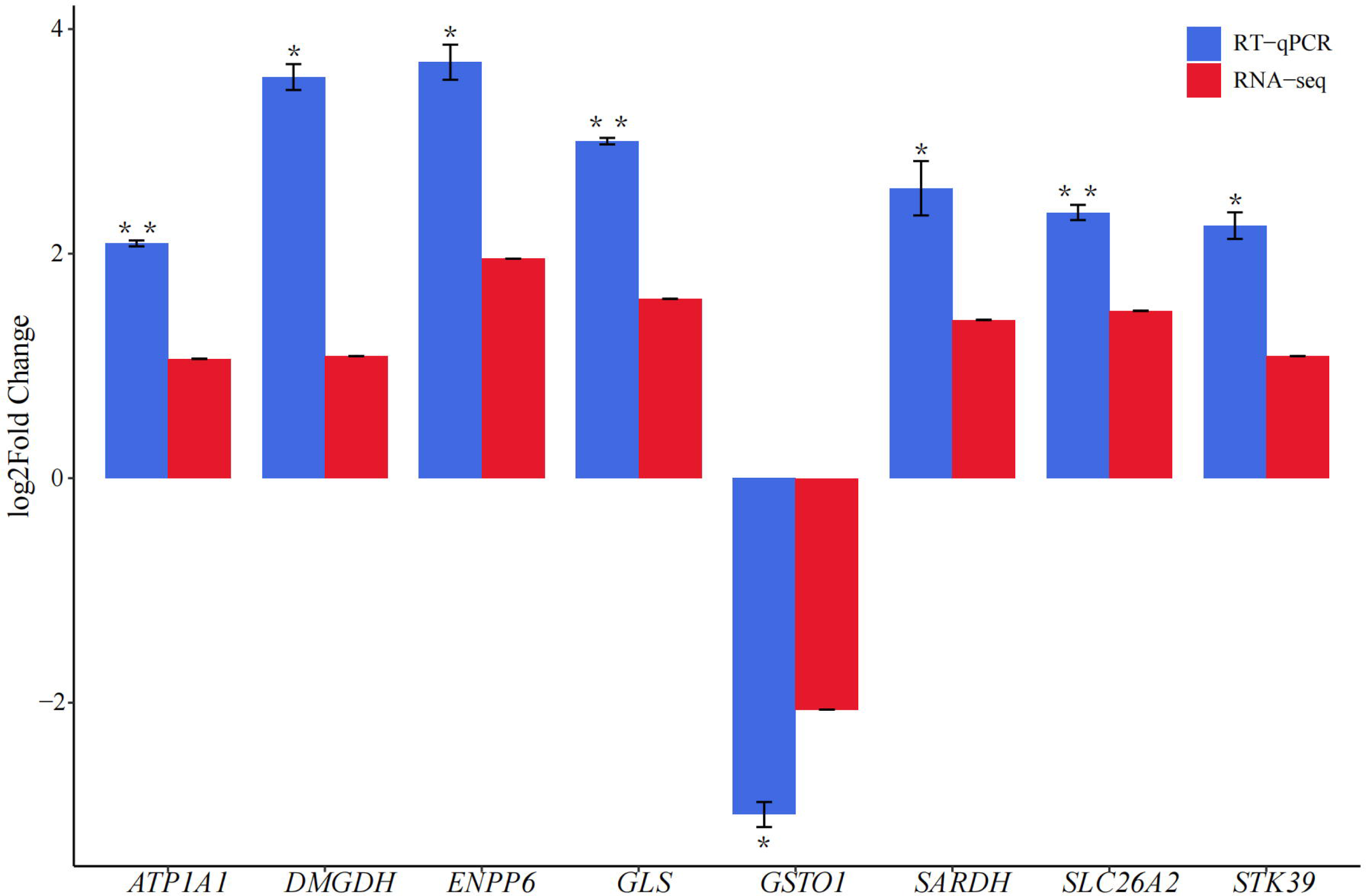
RT-qPCR verification of 8 differentially expressed genes. Between groups of *F. chinensis* in 10 ppt (FC10) and 25 ppt (FC25) salinity environments, 7 genes were upregulated (log2 FC ≥ 1) in shrimp from the FC10 group compared to those from the FC25 group, one gene was downregulated (log2 FC ≤ −1). “*” denotes genes with p-value less than 0.05, “**” represents p-value less than 0.01.

## 4. Discussion

### 4.1 Ion-transport Regulation

During in response to low salinity stress, *F. chinensis* have been shown to be able to consume more energy during osmoregulation [6, 8], and they will active the immune-related enzymes, antioxidant system, especially the in hepatopancreas tissue [3–4]. This study reveals the intricate molecular mechanisms employed by Chinese shrimp in response to low salinity stress, encompassing key genes associated with ion-transport regulation, bio-metabolism and detoxification. Ion-transport regulation is an important way for crustaceans to cope with low-salinity stress rapidly, and strategies based on it are known as the “complementary process” of osmoregulation [33]. In the process, Na^+^/K^+^-ATPase, accounting for more than 70% of the activity of ATPase, is the most important example of ion transport enzymes in maintaining the body’s sodium-potassium balance and regulating the lymph osmotic pressure in crustaceans including Chinese shrimp [34]. Upregulation of *ATP1A1* in shrimp exposed to low-salinity condition in the study indicated that they experienced a positive stress response. In short, it is now most generally accepted as the “pump” in osmoregulation has been well reported in previous studies.

Due to diffusive effluxes of cations, anions from the inside of branchial cells will be attracted to the outside of the cells. The branchial epithelial cells of crustaceans have strong permeability for Cl^-^, and a large number of Cl^-^ flow outside into the hemolymph occurring at the basolateral membrane. In order to maintain the intracellular balance of Cl^-^, selective channels were activated [35]. In the study, upregulation of *SCL26A2* should be closed related to the anion (Cl^-^) regulation, as the production of *SLC26A2* appears to function as a SO_4_^2-^/Cl^-^ exchanger, which could move Cl^-^ into cells [36]. According to the model of ion transport into epithelial cells across the apical membrane mentioned by Pequeux (1995), upregulation of *SLC26A2* in this study implied that this gene’s production should be located in the apical membrane providing a channel for the chloride ion (Cl^-^) entering the cell, and by which Chinese shrimp could maintain intracellular chloride balance. In aspects of osmoregulation mechanisms in crustacean, the function of *SLC26A2* may be a supplement point to the mechanism HCO_3_^-^/Cl^-^ - exchange.

Otherwise, as to known, the Ste20 kinase is essential for acute volume recovery and survival after hypertonic shrinkage in *Caenorhabditis elegans*, the founding member of serine/threonine kinases, regulate a *Caenorhabditis elegans* ClC anion channel [37]. In this study, *STK39* as the member of serine/threonine kinases, whose upregulation in response to low salinity stress in Chinese shrimp may seem to make sense. As to date, however, there is no report on the research of the gene’s participation in osmoregulation of shrimp species, thus it is necessary to pay more attention on this gene in the future.

### 4.2 Ion-transport Regulation

The upregulation of *ENPP6* and *GLS* sheds light on the importance of bio-metabolic processes in Chinese shrimp adaptation to low-salinity stress. *ENPP6*, known for its hydrolytic efficiency of glycerophosphorylcholine (GPC), may play a role in reducing organic osmolytes like GPC, aiding the shrimp in adapting to hypoosmotic environments [38]. It was mentioned that in the renal medulla of mammals, GPC plays an important role as an organic osmolyte to maintain intracellular osmotic pressure, and accumulating of organic osmolytes, cells can adapt to environmental hypertonicity by high osmolyte concentrations[39]. In other words, reduction of organic osmolytes in cells such as the GPC may imply Chinese shrimp’s adaptation to hypoosmotic environments, and the involvement of *ENPP6* in the response to low salinity stress highlights the significance of bio-metabolic processes in Chinese shrimp adaptation. However, there are no reports on the gene in relation to shrimp at present. Moreover, the upregulation of *GLS* (glutaminase) in response to low-salinity conditions further emphasizes the importance of bio-metabolic pathways in aspects of the energy supplement. Glutaminase catalyzes the conversion of glutamine to glutamate, playing a key role in amino acid metabolism as well as in energy production[40–41]. This finding suggests a potential shift in energy metabolism to facilitate the adaptation of Chinese shrimp to low-salinity stress. Thus, the modulation of amino acid metabolism might contribute to the synthesis of osmoregulatory molecules or act as a compensatory mechanism for energy production under altered environmental conditions.

Meanwhile, the upregulation of *DMGDH* suggests its involvement in bio-metabolic adjustments, which encodes dimethylglycine dehydrogenase involving in the metabolism of dimethylglycine[42]. It was reported that *DMGDH* expression suppresses human hepatocellular carcinoma metastasis in vitro and in vivo [43]. The role of *DMGDH* in low-salinity adaptation in organism is not well-established, and further research is needed to elucidate its specific contributions. Nevertheless, the observed upregulation suggests its potential involvement in bio-metabolic adjustments to cope with the challenges imposed by low salinity.

### 4.3 Detoxification

The down-regulated expression of *GSTO1* (glutathione S-transferase omega-1) in the low-salinity treatment group raises questions about its role in detoxification processes. *GSTO1* is known for its participation in the detoxification of reactive oxygen species, facilitating the binding of glutathione to oxidized substances and aiding in the efficient elimination of harmful compounds from the body [44–45]. The reduced expression of *GSTO1* in response to low salinity might indicate a lower demand for detoxification pathways under the condition (4 days domestication in 10 ppt water). Alternatively, it could suggest a potential compromise in the detoxification capacity of Chinese shrimp, which warrants further investigation, although it was reported that *GST* is also differently expressed in hepatopancreas of Chinese shrimp in response to low salinity stress [3].

In addition to *GSTO1*, it is noteworthy to different expression of the *CTH* gene. *CTH*, encoding cystathionine gamma-lyase, is a key enzyme in the sulfur metabolism pathway, involved in biosynthesis of cysteine [46]. The negative correlation of *CTH* with the majority of differentially expressed genes (DEGs) implies that the expression of *CTH* is inversely related to genes associated with ion transport, bio-metabolism, and detoxification. Cysteine is an essential amino acid and a precursor to glutathione, a crucial antioxidant molecule. Therefore, like *GSTO1*, the downregulation of *CTH* may negatively impact antioxidant defense mechanisms, contributing to the overall adaptive response to low-salinity stress. This observation is consistent with a previous study and reinforces the importance of examining multiple genes and pathways to gain a comprehensive understanding of the molecular responses to environmental stressors in Chinese shrimp [47].

The different expression of sarcosine dehydrogenase (*SARDH*) under low-salt conditions also suggests its potential role in the adaptive response of shrimp to environmental challenges. Sarcosine can be degraded to glycine by sarcosine dehydrogenase [48]. Glycine is a significant amino acid and participates in the synthesis of glutathione, creatine and uric acid, and serves as an advance signal that activates thyroid function coincident with energy and nutrient intake [4]. As we mentioned above, reduced expression of *GSTO1* in response to low salinity might indicate a lower demand for detoxification pathways, thus the integrated analysis illustrated that upregulation of *SARDH* may play a limited role in detoxification mechanisms in gill tissue.

## 5. Conclusions

In conclusion, our study provides a comprehensive analysis of the molecular mechanisms employed by Chinese shrimp in response to low salinity stress. Through RNA-seq and functional enrichment analyses, we identified key genes associated with ion-transport regulation, bio-metabolism, and detoxification, shedding light on the intricate adaptive responses of Chinese shrimp to environmental challenges. The upregulation of *ATP1A1*, *SLC26A2*, and *STK39* suggests a crucial role of ion-transport regulation in rapidly coping with low-salinity stress, highlighting the significance of Na^+^/K^+^-ATPase and chloride ion regulation in maintaining osmotic balance. Furthermore, the involvement of *ENPP6*, *GLS*, and *DMGDH* in bio-metabolic processes underscores the importance of energy metabolism adjustments for shrimp adaptation to hypoosmotic environments. The differential expression of *GSTO1* and *CTH* raises questions about the nuanced interplay of detoxification pathways, suggesting potential compromises and adaptations in response to low salinity. Our findings contribute to a deeper understanding of the molecular responses of Chinese shrimp to low salinity stress, providing valuable insights for aquaculture practices and further research. Future investigations should delve into the specific roles of identified genes, their interactions, and the broader implications for the adaptive capacity of Chinese shrimp in fluctuating environmental conditions. Overall, this study lays the foundation for exploring targeted interventions and strategies to enhance the resilience of shrimp aquaculture in the face of changing salinity levels.

## Supplementary Materials

Supplement 1: Functional annotation of genes; Supplement 2: Results of difference on gene expression; Supplement 3: Results of GO enrichment analysis.

## Author Contributions

Daizhen Zhang and Jun Liu were pivotal in conceptualizing and designing the study. Rongchen Liu and Hui Jiang took the lead in analyzing the sequencing results, providing critical insights into the data interpretation. Hai Zhang and Dianyuan Zheng were instrumental in conducting the experiments related to the project. Both Rongchen Liu and Jun Liu contributed significantly to the writing and drafting of the manuscript.

## Acknowledgments

We thank Dan Liu, Zhen Wang for their assistance in our work. We thank Dachuan Dong for its linguistic assistance during the preparation of this manuscript. Reviewers provided constructive recommendations for improving our manuscript.

## Data Availability Statement

The datasets utilized in this research endeavor are publicly accessible through online repositories. The datasets can be accessed via the following link: https://www.ncbi.nlm.nih.gov/sra/PRJNA734726.

